# Exploring Harmala Alkaloids as Novel Antimalarial Agents against *Plasmodium falciparum* through Bioinformatics Approaches

**DOI:** 10.1101/2024.07.17.603828

**Authors:** Kaushik Zaman Dipto, Raiyan Shariar, Chinmoy Kumar Saha, Abir Huzaifa, Tanjin Barketullah Robin, Rajesh B. Patil, Md. Tamzidul Alam, Md. Irfan Habib Rafi, Ashraf Zaman Faruk, Abu Tayab Moin, Kazi Md. Ali Zinnah, Md. Hasanuzzaman, Tofazzal Islam

## Abstract

Malaria, caused by the *Plasmodium falciparum*, remains a significant global health challenge, with Plasmodium falciparum accounting for approximately 50% of cases and posing a considerable threat. Despite advances in control measures, malaria continues to cause an estimated one million deaths annually. The complex lifecycle of *P. falciparum*, involving both vertebrate hosts and Anopheles mosquitoes, complicates eradication efforts. The parasite’s resistance to existing antimalarial drugs, along with medication toxicity, necessitates innovative therapeutic approaches.

Recent research has revealed that harmine, an alkaloid produced by an endophytic gut bacterium of Anopheles mosquitoes, can impede the transmission of the malarial parasite to humans by inhibiting a crucial life stage. This study investigates harmala alkaloids, sourced from plants and bacteria such as *Peganum harmala*, as potential alternatives to conventional antimalarial drugs. Notably, harmine and harmaline have shown promising antimalarial activity by inhibiting the essential enzyme protein kinase 4 (PK4), which is vital for the parasite’s survival. These compounds exhibit lower toxicity, effectively inhibiting both the blood stage growth and transmission of the parasite. Using in silico methodologies, including ADME analysis, molecular docking, MD simulation, and toxicity analysis, this study identifies harmala alkaloids as potential inhibitors against crucial *P. falciparum* proteins. Targeting proteins essential for the parasite’s survival, similar to established drugs like pfCRT protein, lays the foundation for developing effective antimalarial treatments. The comprehensive screening of harmala alkaloid molecules opens avenues for the pharmaceutical industry to tackle challenges related to drug resistance and toxicity, offering a promising route for the biorational management of malaria.

## 1. Introduction

Malaria, a longstanding and lethal disease, has afflicted humanity for centuries. Various *Plasmodium* species, such as *P. vivax, P. malariae*, and particularly *P. falciparum*, contribute to distinct types of malaria. *P. falciparum,* known for its severity, can lead to malignant malaria and is responsible for approximately 50% of all reported cases [1]. Globally, malaria is a direct cause of an estimated one million deaths annually [2], contributing nearly 3% to the world’s Disability-Adjusted Life Years (DALYs) [3]. An estimated 550 million individuals are thought to be at risk of contracting malaria each year, which results in one million deaths. According to reports, over 49% of the world’s population resides in regions where malaria is spread, which include 109 nations in sections of Africa, Asia, the Middle East, Eastern Europe, Central and South America, the Caribbean, and Oceania. One thousand seven hundred cases of malaria were reported to the Centers for Disease Control. Of these, 65% were obtained in Africa, 19% in Asia, 15% in the Americas and the Caribbean, and less than 1% in Oceania in 2010. While Africa bears the brunt of global malaria mortality, Southeast Asia, including Bangladesh, grapples with substantial morbidity and mortality, posing a significant public health challenge. [4, 5] The malaria parasite is a type of blood parasite that is transferred by female *Anopheles* mosquito. It produces Sporozoites (Sporozoa). Malaria parasites have a complex lifecycle, involving asexual replication in the liver and blood of vertebrate hosts, as well as in the hemocoel of insect vectors. Additionally, they have to go through one round of sexual reproduction, which happens in the midgut of the vector after it consumes blood.

*P. falciparum* is a unicellular protozoan. It belongs to the Sporozoa class. The female Anopheles mosquito is the only vector of *P. falciparum*, as it is the only mosquito that can harbor the parasite in its salivary glands and inject it into the human blood during a bite. it needs two different hosts to perform asexual and sexual reproduction. Sexual reproduction occurs in the midgut of female Anopheles mosquitoes and asexual development happens in humans (infectious form). During the asexual stage in human, it initially multiplies within the liver cells and then attack the red blood cells (RBCs) resulting in their rupture. The rupture of RBCs is associated with the release of a toxic substance, haemozoin which is responsible for the chill and high fever recurring every three to four days. Similar to other forms of malaria, *P. falciparum* infection can cause more severe and potentially fatal symptoms. Fever (which can reach 41°C or 105.8°F), headache, chills, exhaustion, seizures, breathing difficulties, dark or bloody urine, jaundice, and irregular bleeding are a few signs of *P. falciparum* malaria[6]. The symptoms of a *P. falciparum* infection often appear 10 to 15 days after the mosquito bite, although they can potentially appear weeks or months later if preventive medication is taken or if the liver is in a dormant stage[7].

Some common treatment strategies against *Plasmodium falciparum* are followed. Among them, Antimalarial drug therapy is notable which includes quinine and quinidine, and antifolate drugs[8, 9]. Limitations of current treatment strategies decline the effectiveness of medications.

*P. falciparum* has developed resistance to many antimalarial drugs, including artemisinin-based therapies, making them less effective in some regions [10]. Certain antimalarial medications, such as mefloquine and quinine, can be extremely toxic and have serious side effects. This restricts their use, particularly in those that are more vulnerable[11]. Newer strategies are being developed to effectively cure the deadly effects of *Plasmodium falciparum*. Harmala Alkaloids are proposed as Natural Alternatives to present anti-malarial drugs.

Harmala alkaloids are a group of natural compounds that are derived mostly from plants of the family Zygophyllaceae, such as *Peganum harmala*, and have been studied for their potential antimalarial properties. These compounds, particularly harmine and harmaline, have shown activity against *P. falciparum* in laboratory studies. These compounds show potential novel mechanisms. An enzyme called protein kinase 4 (PK4), vital for the parasite’s survival, is inhibited by them. Recently, harmine alkaloid produced by a gut bacterium of some *Anopheles* mosquitoes was found to inhibit the transmission malarial parasite from mosquito to humans through impairing a particular life stage of the *P. falciparum*. This provides a unique treatment for malaria and could help to overcome drug resistance [12]. These compounds have less toxicity compared to other antimalarial drugs available and can inhibit the growth of the parasite in the blood stage, as well as prevent its transmission by reducing the production of gametocytes, the sexual stage that infects mosquitoes.

While the antimalarial effect of harmine produced by a gut bacterium of Anopheles mosquitoes has been demonstrated [13], the structure-activity relationships of harmala alkaloids and their comparative efficacies on the target proteins of the parasite remain unclear. In this study, we focused on essential proteins targeted by established drugs like the pfCRT protein, as detailed in the supplementary information, which play a critical role in the survival and infection of *P. falciparum*. Employing an in-silico approach, we systematically screened potential drug candidates against the malarial parasite. The virtual screening procedure included ADME analysis, molecular docking, MD simulation, MMGBSA, and toxicity analysis. Our results suggest that harmala alkaloids, as natural inhibitors, show promise as candidates for new drugs against deadly malaria.

## 2. Methods

The stepwise methods of the whole study is illustrated in **Figure 1**.

**Figure 1.**
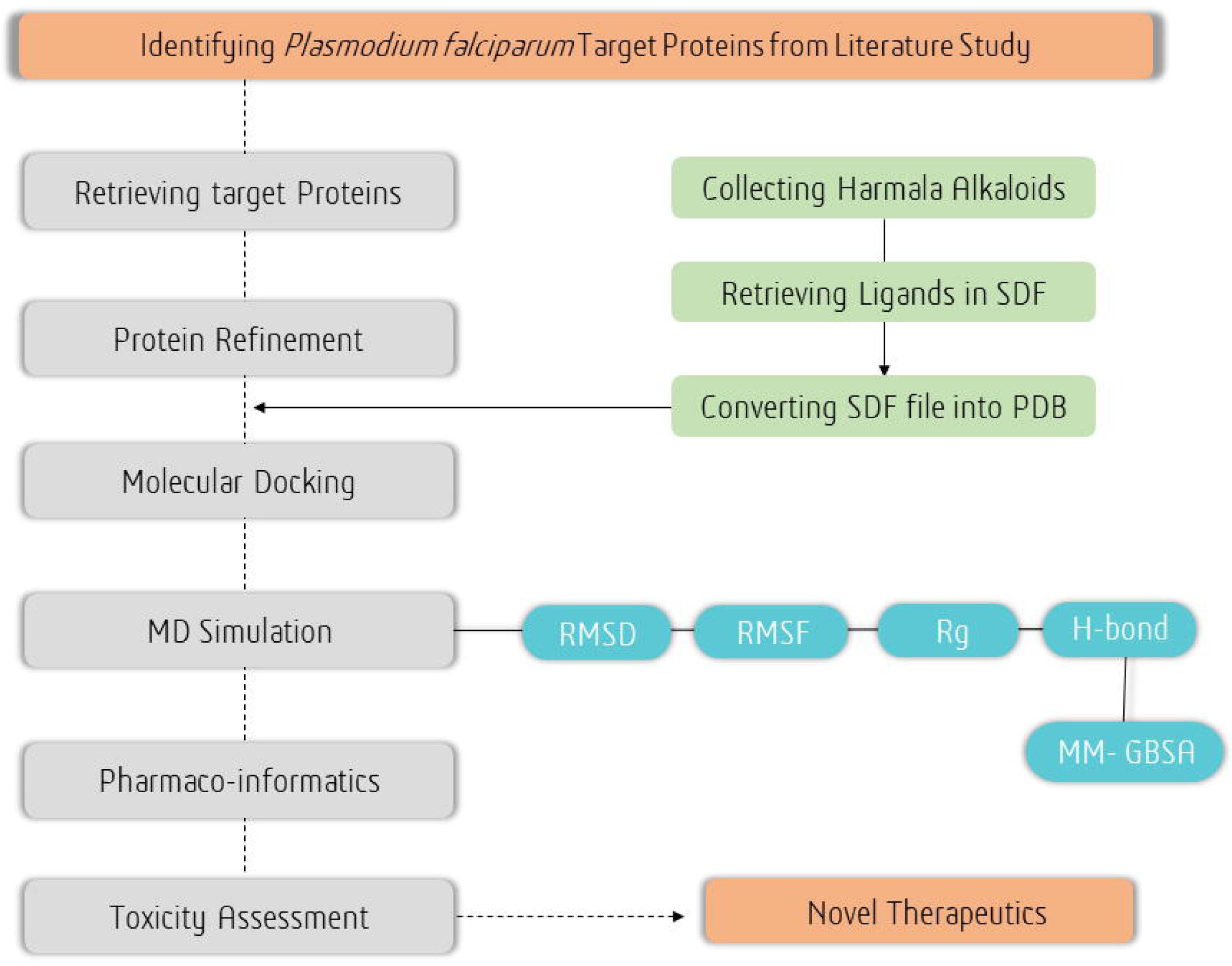
Flowchart depicting the stepwise methods followed in the whole study.

### 2.1. Target protein Selection

Five protein targets were selected after an extensive literature review. The target proteins play a crucial role in *Plasmodium falciparum* survival and infection as given in **Supplementary Table S1**

### 2.2. Pre-Docking Preparation of Protein

The BIOVIA Discovery Studio Visualizer tool was used to refine the target protein by removing unwanted ligands, metals, and ions [14]. The program aids in the optimization of compound selection by enabling the visualization, profiling, and analysis of chemical libraries from various sources.

### 2.3. Collecting Potential Inhibitors

We carefully selected 21 harmala alkaloid metabolites having potent anti-protozoal and anti- microbial activity were selected after the literature review. The source of the selection and their importance is given in (**Supplementary Table S2**). These compounds, metabolites, and inhibitors exhibit promise as therapeutic possibilities for treating malaria. The two most widely used malarial drugs Quinine and Chloroquine were used as control to check the potential of novel metabolites [8, 15].

### 2.4. Ligand Preparation

The PubChem database was used to retrieve the metabolite structures, which were provided in SDF (3D) format (https://pubchem.ncbi.nlm.nih.gov/)[16]. The flexible chemical data programmed Open Babel v2.3, which supports over 110 distinct formats, was used to convert the structures to PDB format in order to prepare them for additional investigation [17].

### 2.5. Molecular Docking Analysis

Molecular docking is an important method for studying interactions between small ligands and large molecules with minor changes. As a result, the Python Prescription 0.8 (PyRx) package’s Auto Dock Vina was used to examine the molecular docking of the selected ligands with our target proteins [18]. The protein data bank, partial charge, and atom type (PDBQT) files were built using the protein’s previously constructed PDB files as inputs. All 21 compounds and two reference medicines were minimized and transferred to PDBQT format before docking. Vina Wizard was used to do the docking procedure. The greatest size of the grid for the receptor was chosen for the investigation, with the grid’s center at X: 23.171, Y: 20.8184, and Z: 7.2116. The points were X:164, Y:86, and Z:121, and the space had a value of 0.3750Å.

### 2.6. Binding site and complex evaluation

The PyMOL v2.0 program and the BIOVIA Discovery Studio were used to examine and visualize binding locations. PyMOL, a popular program, can be used to visualize biological macromolecules like proteins and tiny chemicals in 3D. It is extremely useful for displaying ligand-binding complexes. PyMOL, a proprietary, open-source molecular visualization tool created by Warren Lyford DeLano, is one of the most important open-source tools for visualizing structural biology. PyMOL is used to identify polar and non-polar residues and to investigate where specific metabolites bind [19, 20]. Discovery Studio was used to see if any unintended interactions occurred during the investigation.

### 2.7. Molecular Dynamics Simulation studies

The docking results suggested promising binding free energies of Harman-3-carboxylic acid (HC) and Harmine N-oxide (HO) compared to the standard antimalarial drug chloroquine (CQ) for the targets AMA1, MSP1, pfCRT,pfEMP1 and pfPK5. To have insights into the binding free energies in a simulated biologically relevant environment, the complexes of these ligands with the targets were subjected to molecular dynamics simulations. The 100 ns MD simulations were performed using the Gromacs-2020.4 [21, 22] program using the resources of the HPC cluster at Bioinformatics Resources and Applications Facility (BRAF), C-DAC, Pune. The CHARMM-36 force field parameters [23, 24] were employed while generating the topologies of respective proteins, and the ligand topologies were obtained from the CGenFF server [24, 25] The complexes were solvated using the TIP3P water model [26] in a dodecahedron unit cell, maintaining a distance of 1 nm between the unit cells and the complexes’ boundaries. The resulting solvated systems were neutralized by sodium and chloride ions, maintaining the molar concentration of 0.15 mole. The stearic clashes in the system were relieved by the steepest descent energy minimization, where the threshold of the force constant was set to 100 kJ mol^-1^ nm^-1^. Subsequently, two-step equilibrations of 1 ns each at constant temperature (NVT) and constant pressure (NPT) were performed on each system. During NPT equilibration the constant temperature of 300 K was achieved with a modified Berendsen thermostat [27]; while during NPT equilibration the constant pressure of 1 atm was achieved with a Berendsen barostat [28]. However, during the 100 ns production phase MD simulations, the pressure conditions of 1 atm were achieved with the Parrinello-Rahman barostat [29], while the temperature of 300 K was achieved with a modified Berendensen thermostat. The position restraints on covalent bonds were achieved with the LINCS algorithm[30]. The long-range electrostatic energies were computed with the Particle Mesh Ewald (PME) method [31] with a cut-off of 1.2 nm. The periodic boundary conditions were removed from the final trajectories obtained from the MD simulations. The trajectories were analyzed for the root mean square deviations (RMSD) in the backbone atoms of proteins, the root mean square fluctuation (RMSF) in the side chain atoms, the radius of gyration (Rg), and the contact frequency between residues and the ligands. The results of contact frequency were compared with the hydrogen bonds between the binding site residue and ligand. The major path of motions in each complex was analyzed from principal component analysis (PCA) [32]. The covariance matrix for the protein backbone atoms was constructed with the gmx cover program. The covariance matrix was diagonalized to obtain the eigenvectors and eigenvalues, where the eigenvectors represented the path of motion, while eigenvalues represented the mean square fluctuations. The first two principal components (PC1 and PC2) were further used to obtain Gibb’s free energy landscape [33]. Molecular mechanics energies combined with Poisson Boltzmann surface area continuum solvation (MM-PBSA) calculations were performed on the trajectories isolated at every 1 ns from 75 to 100 ns simulation period, employing the GMX_MMPBSA program [34].

### 2.8. ADME Analysis

Drug levels and availability in animal tissues are significantly influenced by their absorption, distribution, metabolism, and excretion (ADME). Studies have shown that early evaluation of ADME during drug discovery can reduce the risk of pharmacokinetic-related failures during clinical stages [35]. In silico methods have been proposed as a reliable alternative to experimental techniques for predicting ADME, particularly during the early stages when the number of compounds to be tested is large but the availability of actual compounds is limited. The SwissADME server was used to assess the ADME characteristics of the most important metabolites [36]. The drugs were uploaded in SDF format and converted to SMILES, and predictions were generated using the server. The BOILED-Egg model was used to determine the blood-brain barrier (BBB) in the compounds of interest [37]. Additionally, the SwissADME tool was employed to quickly establish the bioavailability of the ligands [38].

### 2.9. Toxicity Analysis

The pkCSM web-based system (https://biosig.lab.uq.edu.au/pkcsm/) was used to forecast the general toxicity of the discovered chemicals [39]. This program uses graph-based signatures to assess molecular distance patterns, making it effective for predicting pharmacokinetic aspects.

## 3. Results

### 3.1. Selecting and retrieving the target proteins

The target proteins of the malarial parasite, *P. falciparum* viz. MSP1, pfPK5, pfCRT, pfEMP1 and AMA1 were retrieved from RCSB Protein Databank with PDB ID 1OB1, 1OB3, 6UKJ, 6S8T and 7F9N, respectively.

### 3.2. Protein and ligand preparation

Retrieved proteins were cleaned and unwanted molecules were removed using BIOVIA Discovery Studio software. Then their 3D structures were visualized and given in **Supplementary Figure S1**. A total of 21 Harmala alkaloid metabolites were collected in 3D SDF format from PubchemDatabase. They were further converted into PDB format using open babel software.

### 3.3. Molecular Docking and Binding site analysis

Docking was performed against all 21 metabolites and 2 reference compounds Quinine and Chloroquine (**Supplementary Table S3**). Among all the compounds Harman-3-carboxylic_acid, Harmalol_hydrochloride, Harmalol and Harmine_N-oxide showed significant binding affinity even better than the reference drugs. Among these top candidates Harmine_N-oxide and Harman-3-carboxylic_acid showed the highest binding energy with most of the target proteins (**Table 1**). Quinine (control) showed the lowest energy of -7.3 kcal/mol with pfPK5 protein, which is the best score of our reference drug. Whereas Harmine_N-oxide showed highest binding affinity of -8.3 kcal/mol with pfPK5 protein and a binding site with *LYS32, THR14, VAL18, ASP143, GLN129, LEU132, ASP85, ASP83, LYS88* residues. This novel metabolite also showed the highest binding site (9) which is considered as hotspot of the complex. Here, Quinine and chloroquine showed the highest of 8 binding residues. Their interaction is given in **Supplementary Figure S2-S6.**

**Table 1:**
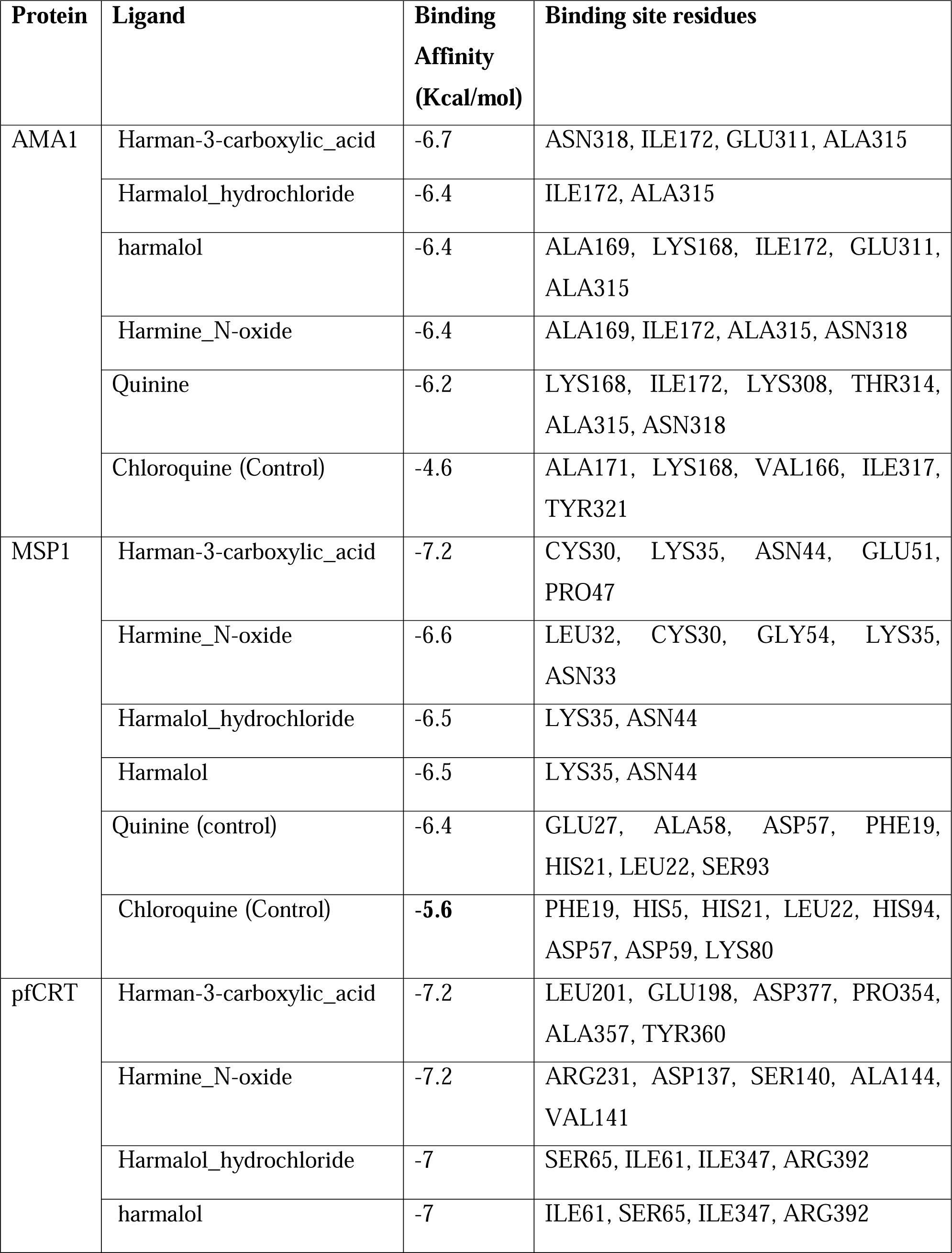
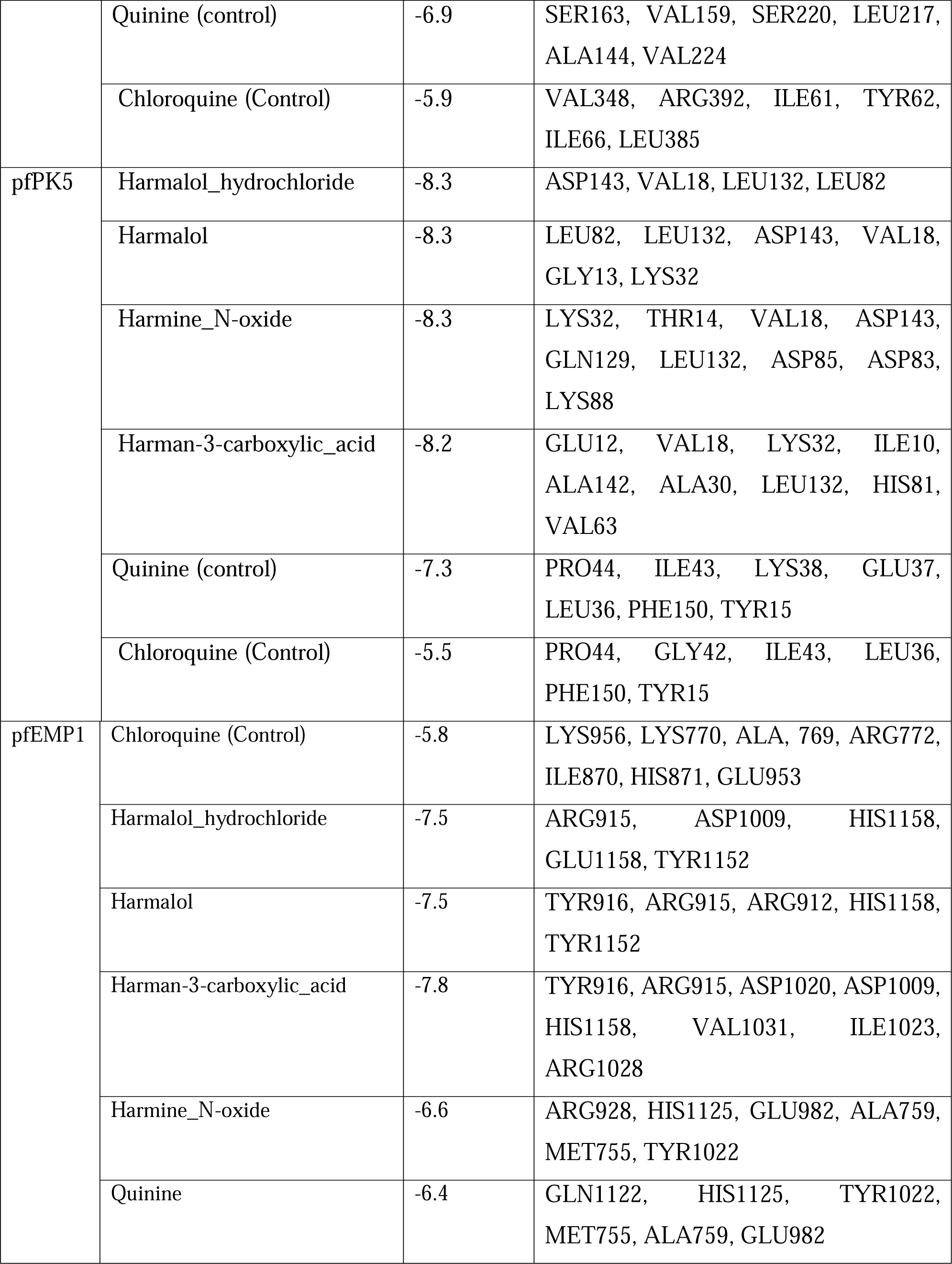
Docking Score and binding sites of top metabolites.

### 3.4. Molecular Dynamics Simulation studies and MM-PBSA calculations

In case of protein *AMP1*, the ligands CQ, HC, and HO were bound at the interface of two chains of AMP1. The MD simulations were performed by taking the complexes of AMP1 with both the chains and ligands. The RMSD, RMSF, and Rg analysis was separately carried out for individual chains of AMP1. The RMSD in AMP1 chain A was stable for complexes with CQ and HO, with an average of around 0.2 nm (**Figure 2**). While considerable deviations were observed in the RMSD in AMP1 chain A until 75 ns, significantly higher deviations were observed until the end of the simulation. The RMSD for AMP1 chain B remained reasonably stable throughout the simulation period for all three complexes. However, the AMP1 complex with HC showed slightly larger deviations from 25 ns to 60 ns during the simulation period. The RMSD in ligand atoms indicated that HO had the least deviations compared to HC and CQ. The RMSD in HC deviates intermittently, but the maximum RMSD remained below 0.1 nm throughout the simulation. The RMSD in CQ showed larger structural changes until 50 ns with an average RMSD of 0.25 nm and stabilizing to a unique conformation after that. The Rg of AMP1 chain A in complex with CQ and HO is reasonably constant throughout the simulation, with an average of around 2.37 nm. The Rg in the HC bound complex showed deviations, which significantly increased after 75 ns. The Rg in AMP1 chain B was similarly observed to be quite stable in the case of complexes with CQ and HO, while the Rg deviated significantly in complexes with HC after around 25 ns simulation period. The RMSF analysis suggested that the fluctuations in side chain atoms of AMP1 chain A were comparably lower in all the complexes. However, the complex with CQ showed significant fluctuations in the side chain atoms of residues in the range 240 to 250, while the complex with HC showed significantly larger fluctuations in the side chain atoms of residues in the range 260 to 275. In the case of AMP1 chain B, the RMSF remained almost the same magnitude for all the complexes and slightly larger fluctuations in the side chain atoms of residues in the ranges 240 to 250 and 260 to 275.

**Figure 2.**
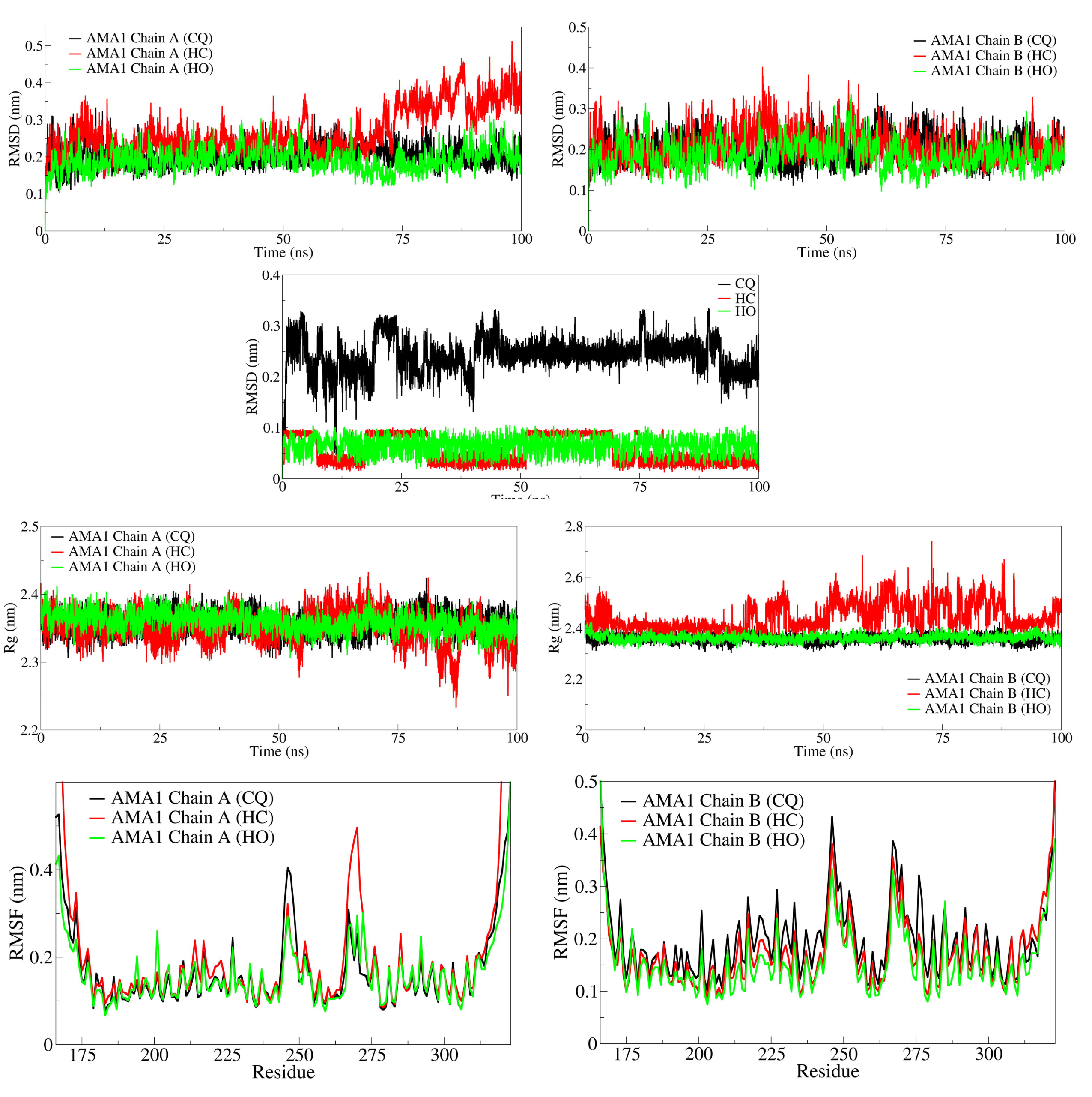
RMSD, Rg, and RMSF analysis for AMP1 complexes.

The hydrogen bond analysis showed that the AMP1 complex with CQ formed a consistent hydrogen bond throughout the simulation period and occasionally formed two hydrogen bonds (**Figure 3**). The contact frequency between ligands and residues within 0.35 nm was analyzed. The contact frequency analysis showed that CQ makes contacts with more than 25% contact frequency with residues Tyr321, Gln313, Val166, Lys168, and Thr320. In the case of the AMP1 HC complex, the number of hydrogen bonds was fewer than the CQ complex. Further, the contact frequency analysis suggested very subtle and weak contacts with residues Glu177, Ala169, Gln303, Ser304, and Asn318 with less than 5% contact frequency. The AMP1 HO complex showed very few hydrogen bonds during the initial simulation and at around 30 ns simulation period. Further, the contact frequency with residues Lys168, Glu311, Ala169, Lys308, and Asn318 was lower than 10%.

**Figure 3.**
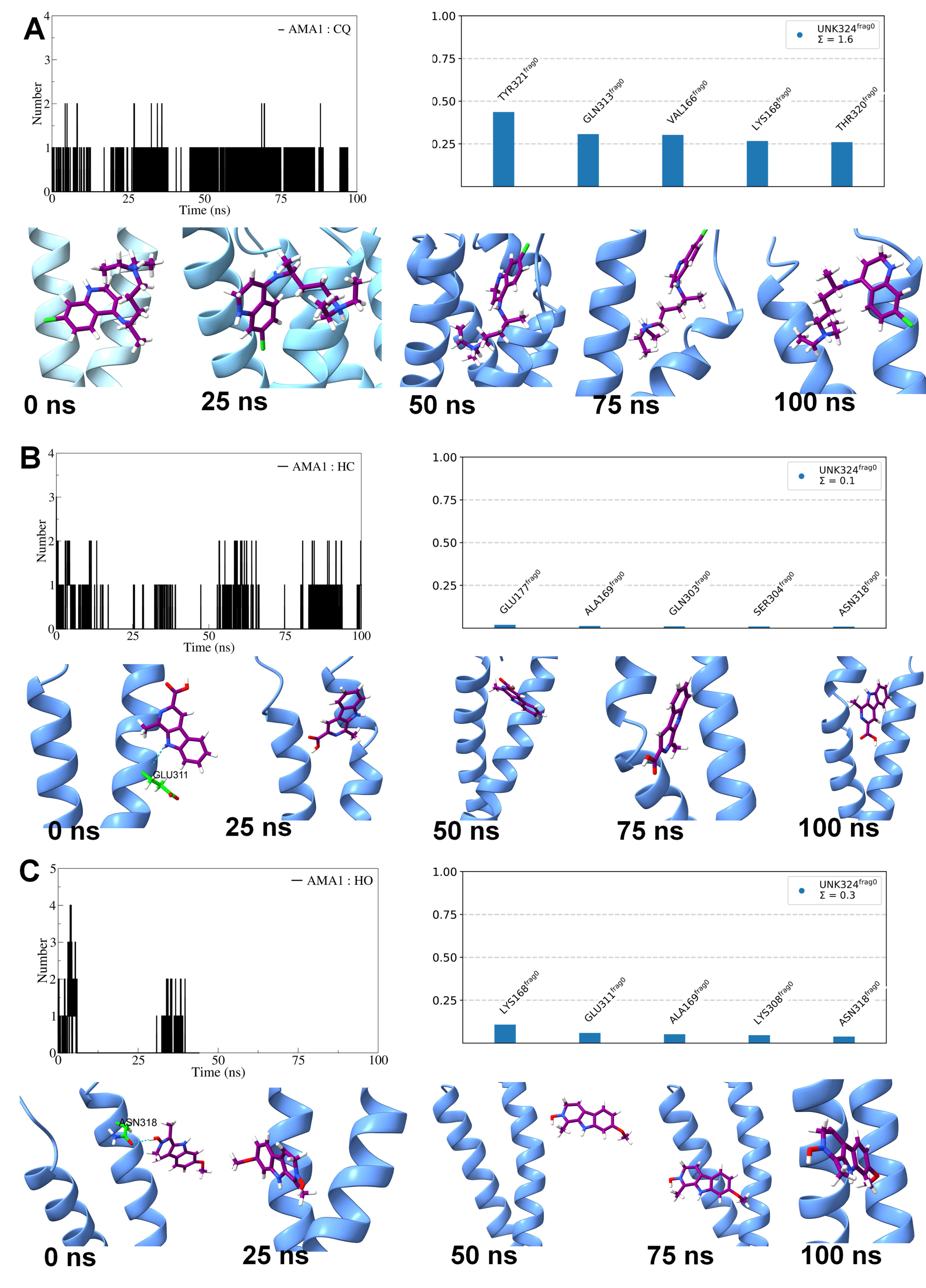
Hydrogen bond and contact frequency analysis for AMP1 complexes.

In case of protein *MSP1* the RMSD in MSP1 backbone atoms in the complex with CQ was the lowest, with an average of around 0.25 nm (**Figure 4**). In the MSP1 complex with HC, the RMSD was found stable with a slightly higher average of around 0.3 nm compared to the MSP1 complex with CQ. The RMSD in backbone atoms of MSP1 in complex with HO deviated significantly until 75 ns and thereafter stabilized to an average of 0.36 nm. The RMSD in ligand atoms suggested that the ligand HC has the least deviations with an average of below 0.05 nm. The ligand HQ has a significantly lower RMSD with an average of 0.08 nm, while the atoms of CQ deviated throughout the simulation with a significantly higher RMSD with an average of 0.2 nm. The RMSF analysis showed that MSP1 in complex with HO had higher fluctuations in the side chain atoms of residues. Particularly, the fluctuations were seen in the residues in the ranges 30-60 and 80-90. Comparably, the RMSF in MSP1 in complexes with CQ and HC was lower and almost similar. The Rg was found stable for all the complexes of MSP1 with an average of around 1.41 nm. However, compared to the MSP1 complex with CQ, the complex with HC and HO showed significantly higher Rg during the initial 15 ns simulation period.

**Figure 4.**
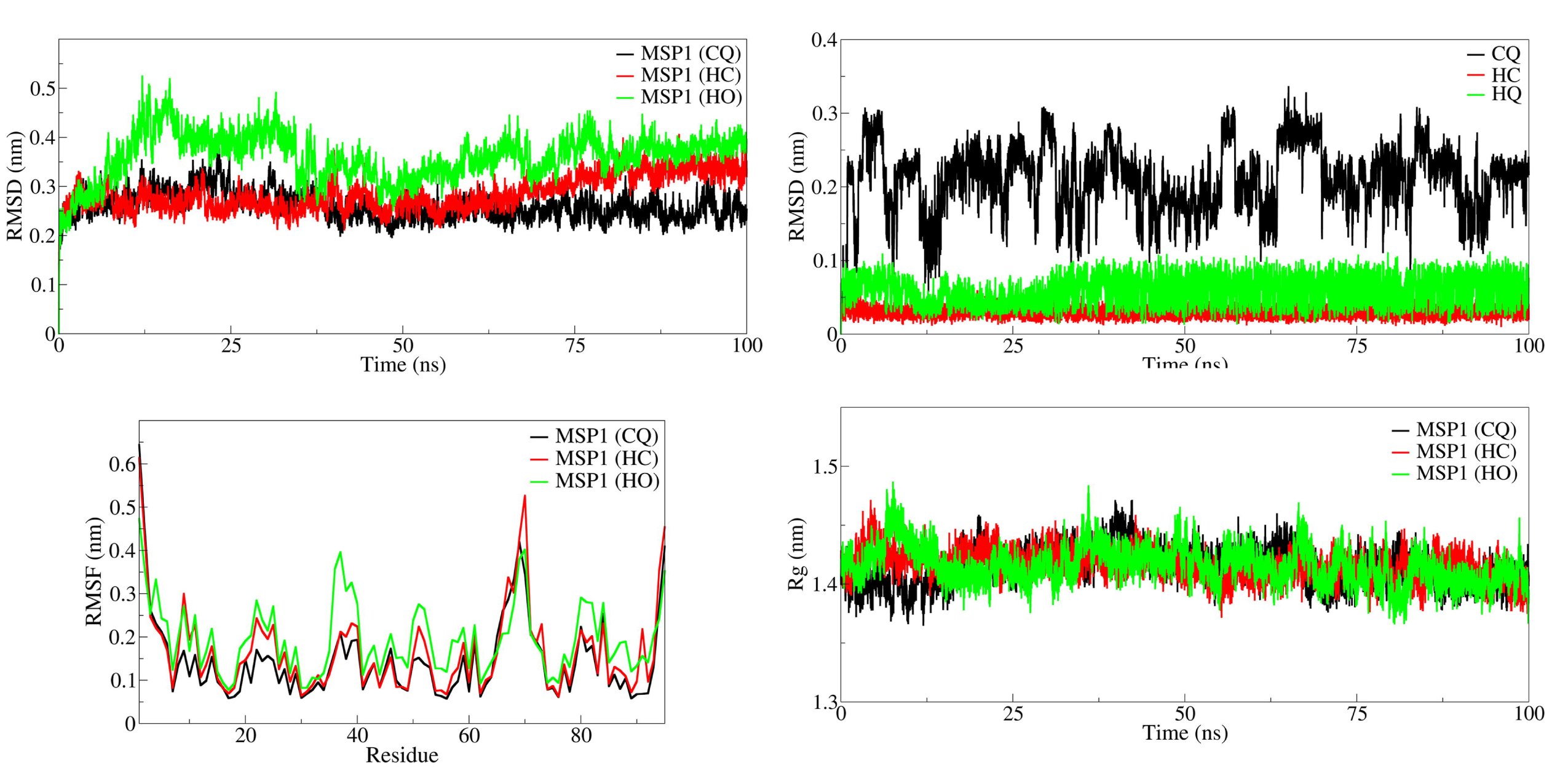
RMSD, Rg, and RMSF analysis for MSP1 complexes.

The hydrogen bond analysis showed that CQ formed significantly less frequent hydrogen bonds with MSP1, while HC formed a significantly higher number, for instance, 4 hydrogen bonds, out of which 3 hydrogen bonds were consistently formed throughout the simulation period (**Figure 5**). However, the ligand HO formed around 3 hydrogen bonds less frequently than HC. The contact frequency for the ligand CQ showed that the residue Lys80 had more than 50% contact frequency, while Asp59 and Asp82 had more than 25%, and residues His94 and His95 had around 10% contact frequency. On the other hand, the ligand HC showed more than 75% contact frequency for the residues Thr48 and Asn44 and more than 50 % contact frequency for the residues Gly54, Asn52, and Gln36. Comparably, the contact frequency for the HO showed a significantly lower contact frequency of around 10% with the residues Gly54, Cys30, Gln36, Tyr34, and Asn52.

**Figure 5.**
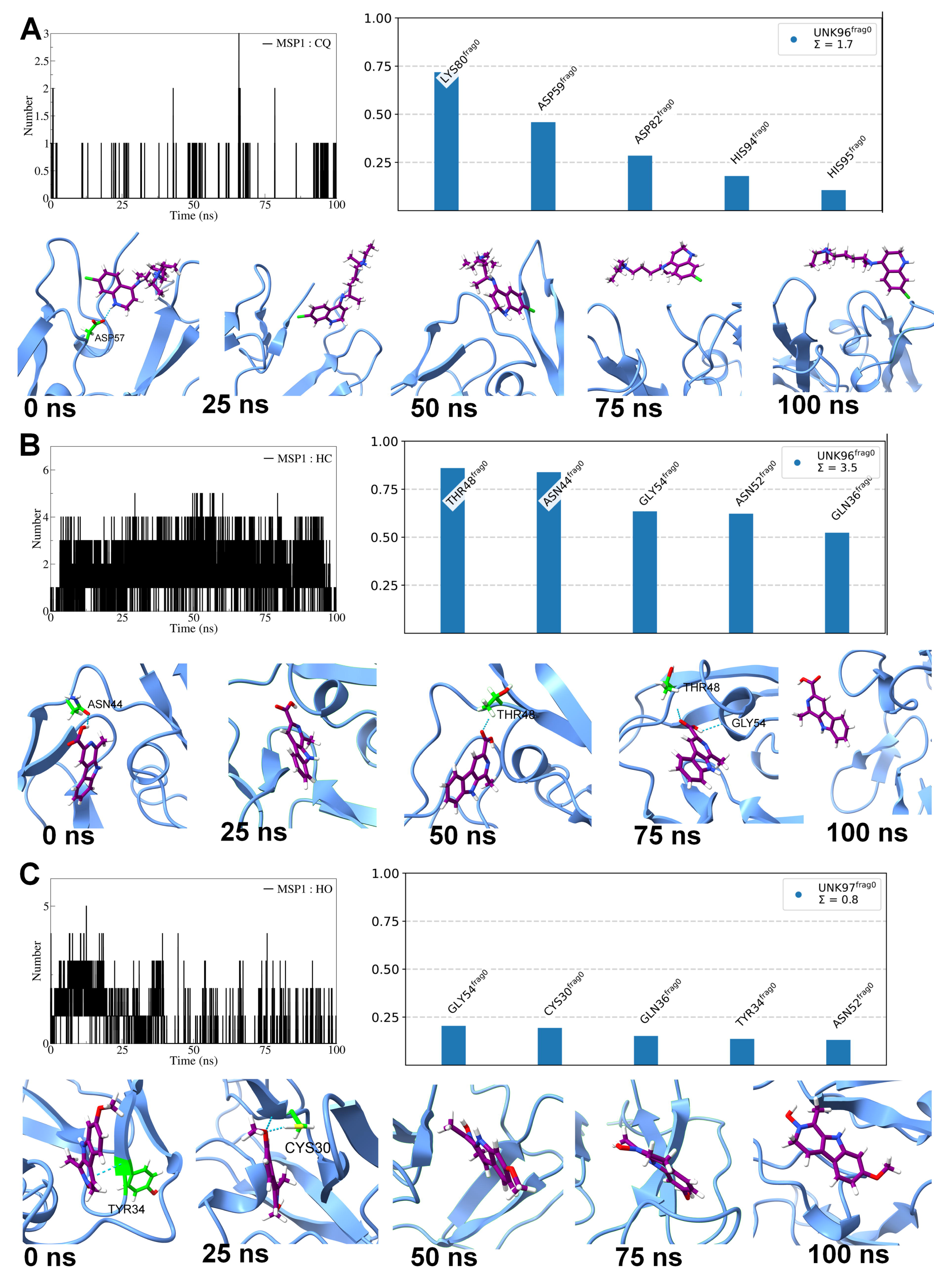
Hydrogen bond and contact frequency analysis for MSP1 complexes.

In case of pfCRT, the RMSD in backbone atoms of the pfCRT complex with CQ was lowest, with an average of around 0.39 nm, whereas the complex with HC was slightly higher, with an average of around 0.4 nm (**Figure 6**). The RMSD in backbone atoms of the pfCRT complex with HO was significantly higher, with an average of around 0.5 nm. The RMSD in ligand atoms showed that the ligands HC and HO had significantly lower RMSD with an average of around 0.09 nm compared to the CQ, for which the RMSD deviated significantly throughout the simulation with an average of around 0.26 nm. The RMSF analysis showed that the fluctuations in the side chain atoms of pfCRT were almost similar in all the complexes. However, the major fluctuations were evident in the residues in the ranges 190-220, 290-310, and 360-380. The Rg analysis showed that the Rg of pfCRT complexes with CQ and HC remained stable throughout the simulation. The Rg of the pfCRT complex with HO was higher, with an average of around 2.23 nm.

**Figure 6.**
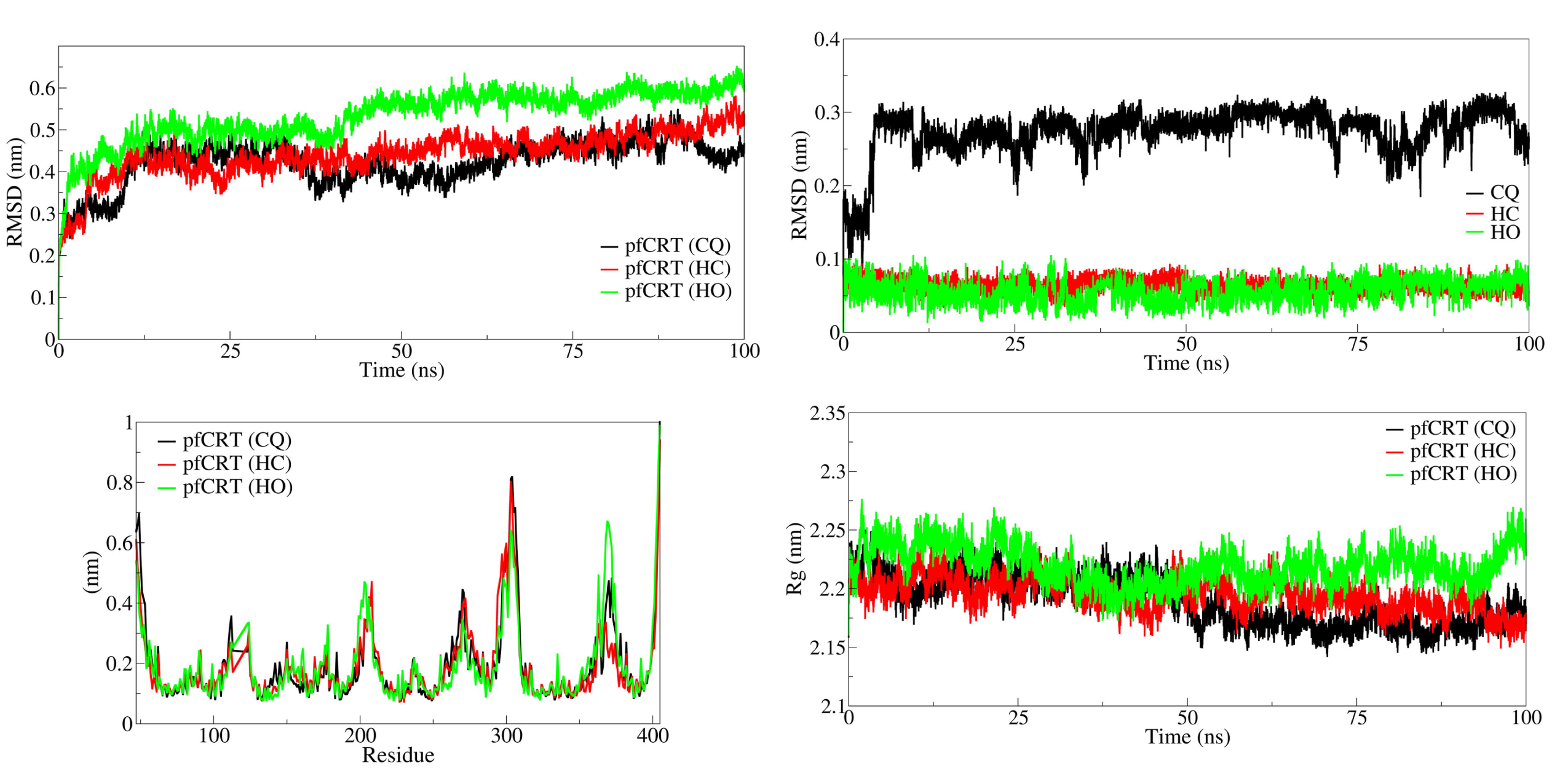
RMSD, Rg, and RMSF analysis for pfCRT complexes

The hydrogen bond analysis showed that CQ formed one consistent hydrogen bond, reaching a maximum of two until around 50 ns and one consistent hydrogen bond after 75 ns until the end of the simulation period (**Figure 7**). The ligand HC formed more consistent hydrogen bonds than CQ, reaching a maximum of 4 hydrogen bonds, and two hydrogen bonds consistently formed throughout the simulation period. The ligand HO formed around 5 hydrogen bonds until the first 50 ns simulation period and 3 hydrogen bonds after 50 ns until the end of the simulation period. The contact frequency analysis showed that the ligand CQ had more than 75% contact frequency for Arg392 and Asn58 and more than 20% for the residues Glu54, Ser388, and Thr344. The ligand HC showed a contact frequency of more than 25% for the residues Asn84 and Gly353, while contact frequency was less than 25% for Ser157, Gln161, and Asp377. The ligand HO showed more than 75% contact frequency for the residues Asn246 and Arg231 and around 50% for Asn330, Gln253, and Asn98.

In case of *pfEMP1,* the RMSD in backbone atoms of pfEMP1 in complex with all the ligands is reasonably stable. However, during the initial simulation period of around 50 ns, the RMSD in backbone atoms of pfEMP1 was lower for complexes with HO compared to complexes with HC and CQ (**Figure 8**). After the 50 ns simulation period, the RMSD remained almost similar for the complexes of HO and CQ, while few deviations were seen in RMSD in the complex of HO. The RMSD in ligand atoms showed that the ligand HO has the lowest RMSD with an average of around 0.005 nm. The RMSD in HC deviated between 0.005 nm and 0.1 nm at various time intervals of simulation. The RMSF in side chain atoms of pfEMP1 remained almost similar for all the complexes except for HC, which showed fluctuations for residues in the ranges 790-800, 950-990, and 1100-1120. The Rg for all the complexes remained almost stable, averaging around 2.35 nm. However, higher deviations in Rg were seen in the pfEMP1 complex with CQ after around 75 ns simulation period.

The hydrogen bond analysis showed that CQ formed around 3 hydrogen bonds with pfEMP1 during the first 10 ns simulation period, stabilizing to 2 hydrogen bonds thereafter until the end of the simulation. Out of these 2 hydrogen bonds, one hydrogen bond was consistently formed throughout the simulation. In the case of HC, the 2 hydrogen bonds are consistently formed throughout the simulation, and occasionally 3 hydrogen bonds were observed. Compared to CQ and HC, HO formed more number of hydrogen bonds. However, during the initial 50 ns simulation period, around 2 hydrogen bonds were formed, and after the 50 ns simulation period, the number of hydrogen bonds formed increased to around 5. Contact frequency analysis showed that the ligand CQ has more than 75% contact frequency with the residue Asp1020, around 50% contact frequency with Arg915 and Glu1156, and around 25% contact frequency with residues Ile1023 and Arg1028. In the case of ligand HC, more than 75% contact frequency was observed with residues Asp1020, Asp1009, and Arg912, while a contact frequency of around 50% was observed with the residues Arg915 and Glu1156. In the case of HO, the residues Glu1156 and Tyr1152 showed a contact frequency of more than 75%, while residues Arg915 and Asp1009 showed a contact frequency of around 40%, and residue Ile1023 a contact frequency of around 10% (**Figure 9**).

In case of *pfPK5,* the RMSD in backbone atoms of pfPK5 in complex with CQ was lowest with an average of around 0.21, whereas in complex with HO, it was reasonably stable and lower with an average of around 0.22 nm. Contrary to this, the pfPK5 complex with HC showed significantly higher RMSD with an average of around 0.6 nm until 75 ns and thereafter with an average of around 0.7, reaching a maximum of 0.9 nm at around 80 ns. The RMSD in ligand atoms showed the lowest and most stable RMSD in HQ, with an average of around 0.05 nm. The RMSD in HC atoms deviated between 0.004 to 0.1 at various time intervals. The RMSD in CQ atoms showed larger deviations with an average of around 0.25 nm (**Figure 10**).

The hydrogen bond analysis showed that CQ formed occasional hydrogen bonds at different intervals. The ligand HC formed more hydrogen bonds than CQ, where around 2 hydrogen bonds were consistently formed. Similarly, the ligand HO also formed more hydrogen bonds, of which 2 hydrogen bonds were consistently formed. The contact frequency analysis showed that CQ had around 50% contact frequency for the residues Lys127, Asp125, and Leu126 and around 5% for the residues Tyr15 and Arg35. The ligand HC showed a significantly lower contact frequency, around 5%, for the residues Ile10, Glu12, Gly13, Val18, and Gly11. Similarly, the ligand HO also showed a significantly lower contact frequency of around 5% for the residues Leu82, Asp85, Asp143, Ile10, and Gln129 (**Figure 11**).

#### Gibbs free energy analysis

Gibb’s free energy analysis showed that the AMP1 complex with CQ had numerous low-energy basins with metastable conformations with coefficients between -30 to 25 on PC1 and -20 to 10 on PC2. The AMP1 complex with HC showed a small energy basin with metastable conformations between coefficients 40 to 50 on PC1 and 40 to 50 on PC2. In the case of the AMP1 complex with HO, the metastable conformations were found in a slightly larger low-energy basin occupying the region with coefficients -40 to -15 on PC1 and -12 to -5 on PC2.

In the case of MSP1, the complex with CQ showed a unique low-energy basin between -8 to -4 on PC1 and -4 to 0 on PC2. The complex with HC showed three slightly larger low-energy basins occupying between -3 to 2 on PC1 and -2 to 2 on PC2. The complex with HO showed very few conformations occupying the small energy basin between -40 to -30 on PC1 and -5 to 0 on PC2.

The complex of pfCRT with CQ showed two lowest energy basins, a larger one occupying between 3 to 6 on PC1 and -2 to 0 on PC2 and a smaller one between -5 to -4 on PC1 and 2 to 4 on PC2. The complex of HC showed two lowest energy basins, out of which the larger energy basin occupied the region between 4 to 6 on PC1 and -2 to 0 on PC2, while the smaller one occupied between -7 to -6 on PC1 and 0 to 2 on PC2. The complex of HO showed a large low energy basin between 2 to 6 on PC1 and -2 to 0 on PC2.

The complex of pfEMP1 with CQ showed two low energy basins, out of which one is significantly larger, occupying the regions 1 to 3 on PC1 and -2 to 2 on PC2. The complex of HC showed a unique large low energy basin between 1 to 3 on PC1 and -2 to 0 on PC2. The pfEMP1 complex with HO showed two large low-energy basins, one between the region 2 to 4 on PC1 and -2 to 2 on PC2 and the other between -4 to -3 on PC1 and -2 to 2 on PC2 (Figure 12).

The complexes of pfPK5 showed only CQ having the lowest energy metastable conformations in two small energy basins, one between 1.5 to 3 on PC1 and -2 to 0 on PC2 and the other on the same coordinates on PC1 and 0 to 2 on PC2. Very few low-energy conformations were seen for HC and HO complexes.

#### MM-PBSA calculations

The results of MM-PBSA calculations are shown in **Table 2**. In the case of AMA1 complexes, CQ has the lowest binding free energy with ΔTOTAL of -22 kcal/mol, compared to HC and HO, which have ΔTOTAL of -6.37 and -5.84 kcal/mol, respectively. Comparably, the van der Waals and electrostatic energy were lowest for the AMA1 complex with CQ. In the case of MSP1 complexes, the complex with CQ has the lowest binding free energy with ΔTOTAL of -8.82 kcal/mol. While the complexes with HC and HO have almost similar binding free energy. The polar solvation energy was lower for the MSP1 complex with CQ. The complexes of pfCRT also showed that the complex with CQ has the lowest binding free energy with ΔTOTAL of -25.36 kcal/mol. The complex of pfCRT with HC showed lower binding free energy with ΔTOTAL of - 14 kcal/mol compared to the binding free energy of complex with HO with ΔTOTAL of -5.77 kcal/mol. The van der Waals and non-polar solvation energies were lower for the pfCRT complex with CQ. In the case of pfEMP1 complexes, the complex with HO showed the lowest binding free energy with ΔTOTAL of -18.13 kcal/mol. The complex with HC has a slightly higher ΔTOTAL of -16.39 kcal/mol, and the complex with CQ has a significantly higher ΔTOTAL of -6.34 kcal/mol. The polar solvation energy was comparably higher, and electrostatic energy was lower for the complex of pfEMP1 with HO. In the case of complexes of pfPK5, the complex with CQ has lower binding free energy with ΔTOTAL of -4.98 kcal/mol, while the complexes with HC and HO have ΔTOTAL of -4.3 and -3.92 kcal/mol, respectively.

**Table 2:** The results of MM-PBSA calculations.

| Complex | Energy component (kcal/mol) |  |  |  |  |  |  |
| --- | --- | --- | --- | --- | --- | --- | --- |
| | ΔVDWAALS | ΔEEL | ΔEPB | ΔENPOLAR | ΔGGAS | ΔGSOLV | ΔTOTAL |
| AMA1 complexes |  |  |  |  |  |  |  |
| CQ | -31.76 | -10.53 | 23.61 | -3.33 | -42.29 | 20.29 | -22 |
| HC | -14.69 | -5.36 | 17.58 | -3.9 | -20.05 | 13.68 | -6.37 |
| HO | -13.29 | -6.3 | 15.37 | -1.62 | -19.59 | 13.75 | -5.84 |
| MSP1 complexes |  |  |  |  |  |  |  |
| CQ | -11.08 | -4.26 | 8.42 | -1.9 | -15.34 | 6.52 | -8.82 |
| HC | -13.11 | -6.01 | 16.39 | -1.69 | -19.12 | 14.7 | -4.42 |
| HO | -12.54 | -2.6 | 13.26 | -2.12 | -15.14 | 11.14 | -4 |
| pfCRT complexes |  |  |  |  |  |  |  |
| CQ | -43.74 | -7.77 | 30.41 | -4.27 | -51.51 | 26.15 | -25.36 |
| HC | -28.21 | -21.29 | 38.44 | -2.95 | -49.50 | 35.49 | -14 |
| HO | -28.35 | -31.61 | 57.18 | -2.99 | -59.96 | 54.19 | -5.77 |
| pfEMP1 complexes |  |  |  |  |  |  |  |
| CQ | -31.30 | -12.41 | 41.13 | -3.77 | -43.71 | 37.37 | -6.34 |
| HC | -25.98 | -27.87 | 40.26 | -2.80 | -53.86 | 37.46 | -16.39 |
| HO | -24.04 | -44.21 | 53.20 | -3.08 | -68.25 | 50.12 | -18.13 |
| pfPK5 complexes |  |  |  |  |  |  |  |
| CQ | -12.73 | -1.5 | 12.41 | -3.16 | -14.23 | 9.25 | -4.98 |
| HC | -11.27 | -2.9 | 12.34 | -2.47 | -14.17 | 9.87 | -4.3 |
| HO | -11.93 | -3.15 | 14.56 | -3.4 | -15.08 | 11.16 | -3.92 |

$\Delta$ VDWAALS: van der Waals energy; $\Delta$ EEL: Electrostatic energies; $\Delta$ EPB: Polar solvation energy; $\Delta$ ENPOLAR: Nonpolar solvation energy; $\Delta$ GGAS = $\Delta$ VDWAALS+ $\Delta$ EEL; $\Delta$ GSOLV = $\Delta$ EPB+ $\Delta$ ENPOLAR; ΔTOTAL = $\Delta$ GSOLV + $\Delta$ GGAS

### 3.5. Pharmacoinformatic analysis and bioavailability radar

The drug-likeness properties are important for screening novel drug. The SwissADME server revealed the pharmacoinformatic properties of the top metabolites. All the top metabolites showed good drug-like properties. Their molecular weight was less than 250 Da, no more than 3 H-bond acceptors and 2 H-bond donors. They all perfectly followed Lipinski’s rule. No violation in the drug-like role was shown in the analysis (**Table 3**). The GI absorption of all the predicted metabolites was shown to be high. They also showed a good solubility in water.

**Table 3.** ADME properties of top metabolites.

| **Molecule** | **Harmalol** | **Harmalol_hydrochloride** | **Harman-3-carboxylic_acid** | **Harmine_N-oxide** |
| --- | --- | --- | --- | --- |
| Formula | C12H12N2O | C12H13ClN2O | C13H10N2O2 | C13H12N2O2 |
| MW | 200.24 | 236.7 | 226.23 | 228.25 |
| #Heavy atoms | 15 | 16 | 17 | 17 |
| #Aromatic heavy atoms | 9 | 9 | 13 | 13 |
| Fraction Csp3 | 0.25 | 0.25 | 0.08 | 0.15 |
| #Rotatable bonds | 0 | 0 | 1 | 1 |
| #H-bond acceptors | 2 | 2 | 3 | 3 |
| #H-bond donors | 2 | 2 | 2 | 1 |
| MR | 64.76 | 71.73 | 65.53 | 66.24 |
| TPSA | 48.38 | 48.38 | 65.98 | 47.28 |
| iLOGP | 1.58 | 0 | 1.25 | 2.43 |
| XLOGP3 | 1.75 | 2.55 | 2.58 | 2.08 |
| WLOGP | 1.86 | 2.66 | 2.72 | 2.74 |
| MLOGP | 1.14 | 1.41 | 0.34 | 1.79 |
| Silicos-IT Log P | 3.61 | 3.61 | 2.87 | 1.95 |
| Consensus Log P | 1.99 | 2.05 | 1.95 | 2.2 |
| ESOL Log S | -2.63 | -3.33 | -3.37 | -3.07 |
| ESOL Solubility (mg/ml) | 4.72E-01 | 1.11E-01 | 9.70E-02 | 1.96E-01 |
| ESOL Solubility (mol/l) | 2.36E-03 | 4.67E-04 | 4.29E-04 | 8.60E-04 |
| ESOL Class | Soluble | Soluble | Soluble | Soluble |
| Ali Log S | -2.38 | -3.21 | -3.61 | -2.7 |
| Ali Solubility (mg/ml) | 8.29E-01 | 1.45E-01 | 5.50E-02 | 4.53E-01 |
| Ali Solubility (mol/l) | 4.14E-03 | 6.12E-04 | 2.43E-04 | 1.98E-03 |
| Ali Class | Soluble | Soluble | Soluble | Soluble |
| Silicos-IT LogSw | -3.96 | -3.96 | -4.36 | -3.71 |
| Silicos-IT Solubility (mg/ml) | 2.18E-02 | 2.57E-02 | 9.99E-03 | 4.46E-02 |
| Silicos-IT Solubility (mol/l) | 1.09E-04 | 1.09E-04 | 4.42E-05 | 1.95E-04 |
| Silicos-IT class | Soluble | Soluble | Moderately soluble | Soluble |
| GI absorption | High | High | High | High |
| BBB permeant | Yes | Yes | Yes | Yes |
| Pgp substrate | Yes | Yes | No | Yes |
| CYP1A2 inhibitor | Yes | Yes | Yes | Yes |
| CYP2C19 inhibitor | No | No | No | No |
| CYP2C9 inhibitor | No | No | No | No |
| CYP2D6 inhibitor | No | No | No | No |
| CYP3A4 inhibitor | No | No | No | No |
| log Kp (cm/s) | -6.28 | -5.93 | -5.85 | -6.22 |
| Lipinski #violations | 0 | 0 | 0 | 0 |
| Ghose #violations | 0 | 0 | 0 | 0 |
| Veber #violations | 0 | 0 | 0 | 0 |
| Egan #violations | 0 | 0 | 0 | 0 |
| Muegge #violations | 0 | 0 | 0 | 0 |
| Bioavailability Score | 0.55 | 0.55 | 0.85 | 0.55 |
| PAINS | 0 | 0 | 0 | 0 |
| Brenk | 0 | 0 | 0 | 1 |
| Leadlikeness #violations | 1 | 1 | 1 | 1 |
| Synthetic Accessibility | 2.51 | 2.58 | 1.89 | 2.12 |

### 3.6. Toxicity Assessment

Among the top 4 drugs Harmine_N-oxide and Harman-3-carboxylic_acid showed negative results for AMES toxicity. Though the *insilico* prediction showed AMES Toxicity for Harmalol_hydrochloride and Harmalol, they were found to be non-hepatotoxic in the further evaluation. All the top metabolites showed negative results in hERG I and II inhibitors indicating the metabolites are good for the heart (Table 4). They also showed no skin sensitization and the lethal dose (LD50) was between 3.614 and 2.4.

**Table 4:** Toxicity assessment of top metabolites.

| Toxicity Parameters | Compound Name |  |  |  |
| --- | --- | --- | --- | --- |
|  | Harmalol_hydrochloride | Harmine_N-oxide | Harmalol | Harman-3-carboxylic_acid |
| AMES Toxicity | Yes | No | Yes | No |
| Max. Tolerated Dose (log mg/kg/day) | -0.64 | 0.322 | -0.616 | 0.545 |
| hERG I inhibitor | No | No | No | No |
| hERG II inhibitors | No | No | No | No |
| Oral Rat Acute Toxicity, LD50 (mol/kg) | 2.441 | 3.614 | 2.42 | 2.9 |
| Oral Rat Chronic Toxicity, LOAEL (log mg/kg_bw/day) | 1.632 | 1.556 | 1.786 | 1.267 |
| Hepatotoxicity | No | No | No | No |
| Skin Sensitisation | No | No | No | No |
| *T.Pyriformis* toxicity | 1.043 | 1.501 | 0.854 | 0.28 |
| Minnow Toxicity (log mM) | 0.603 | 1.367 | 0.921 | 0.551 |

## 4. Discussion

In this study, we carried out an extensive *in silico* analysis to evaluate the potential of naturally occurring harmala alkaloids as antimalarial medication candidates. The results of our study indicate that harmala alkaloids have the potential to act as inhibitors of important *P. falciparum* proteins. While earlier studies have revealed the pharmacological properties of harmala alkaloids, such as their anti-inflammatory, anticancer, antidepressant, and antidiabetic properties [44]. However, this study is the investigation, into the molecular understanding of their anti- parasitic effects. The importance lies in recognizing harmala alkaloids as promising contenders, for creating medications to fight against malaria.

The process of discovering antimalarial drugs in the conventional method is quite intricate since it requires a lot of manual work significant financial resources and takes up a considerable amount of time. However, by enabling the quicker discovery of therapeutic targets, computational tools lead to an advancement in the drug development process [40]. The initial stage of creating medications involves the identification of promising targets, within *Plasmodium falciparum* (a type of protozoan) Essential proteins are thought to be the best targets for antimalarial treatment because there is the maximum chances that most drugs will interact with them [41]. Recent research found that *Plasmodium falciparum* has a total of 5,389 proteins. However out of these 1,626 proteins have been classified as hypothetical proteins [42]. Five protein targets were selected after an extensive literature review as given in supplementary table S1 and these are PfCRT, MSP1, AMA1, pfEMP1 and PfPK5 [43] [44] [45] [46] [47] [48] [49].

The target proteins play a crucial role in *Plasmodium falciparum* survival and infection. We chose 21 harmala alkaloid metabolites that have anti-protozoal and anti-microbial properties and docked them with our target proteins. These metabolites are considered to be more environmentally friendly and less harmful, to the environment compared to produced ones. Quinine and chloroquine, the two most commonly used malaria medications, had a much lower binding affinity than any of these protein-ligand complexes. Based on research using molecular docking, the most promising possibilities are believed to be more potent than popular synthetic medications. Harmine N-oxide and Harman-3-carboxylic acid were the leading contenders, exhibiting the highest binding energies with the majority of target proteins. In our docking investigation, Harman-3-carboxylic acid made contact with four polar sites—ASN318, ILE172, GLU311, and ALA315—when paired with the AMA1 protein. Its binding affinity, which was -6.7 Kcal/mol, was the highest. Subsequently, MSP1 containing Harman-3-carboxylic acid displayed five polar residues and achieved the maximum value of -7.2. Subsequently, Harman-3- carboxylic acid exhibited the highest energy of -7.2 when it came into contact with six polar sites (LEU201, GLU198, ASP377, PRO354, ALA357, and TYR360) using pfCRT. Quinine (control) had the lowest energy of -7.3 with pfPK5 protein, which is our reference drug’s highest rating. The highest binding affinity of -8.3 was demonstrated by Harmine_N-oxide with the pfPK5 protein, and a binding site containing residues of LYS32, THR14, VAL18, ASP143, GLN129, LEU132, ASP85, ASP83, and LYS88 was found.

PfCRT, a member of the drug and metabolite transporter superfamily, is found on the membrane of the parasite’s intra-erythrocytic digestive vacuole. In different result of modified form of chloroquine PfCRT showed closely similar result with neutral chloroquine [50]. CQ showed a binding energy of -4.08 kj/mol and CQH+ and CQH2+ showed a binding energy between -4.15 to -4.77. But in this study the Harmala alkaloids outperformed the binding energy result from the neutral chloroquine.

In the pharmacoinformatic analysis, the ADME properties indicates that all the top four compounds are suitable as drug. They significantly follow the Lipinski rule of five. A key idea in medication development is Lipinski’s Rule of Five, also known as the Rule of Five or Ro5. Molecular weight, lipophilicity (logP), hydrogen bond donors, and hydrogen bond acceptors are the four key features that are highlighted. Based on these standards, a compound is considered more promising as a potential medication candidate if it satisfies the following requirements: it must have a molecular weight of fewer than 500 Daltons, a logP value of less than 5, no more than five donors, and no more than ten acceptors of hydrogen bonds [51]. By identifying compounds that are more likely to be orally bioavailable, less toxic, and therefore more viable for further development, the Rule of Five helps researchers in the early stages of drug discovery, ultimately improving the effectiveness and success of drug discovery processes. Our top candidates follow all the rules (Table 3). The MW is between 236 to 200 Da, the H-bond acceptors are between 2-4 and the H-bond donor is between 1-2. The LogP value is also between 0 to 2.43. The toxicity analysis of these metabolites was also significant. Though only Harmalol_hydrochloride and harmalol showed little adverse AMES toxicity effect all of them showed negative results in hERG I inhibitor and hERG II inhibitors. Because of its significance, the hERG channel is included in the list of biomarkers for cardiac safety. hERG I and hERG II are involved in the cardiac system [52]. Inhibiting this system is considered to be dangerous. Again, All the top candidates showed negative results in hepatotoxicity and Skin Sensitization.

The ligand chloroquine (CQ), Harman-3-Because of its significance, the hERG channel is included in the list of biomarkers for cardiac safety. Carboxylic acid (HC), and Harmine N-oxide showed reasonably good binding free energies for the targets AMA1, MSP1, pfCRT, pfEMP1, and pfPK5. To get deeper insights into the binding modes of these ligands and to study the stabilization of these protein targets in complex with the ligands, 100 ns MD simulations were performed. Typically, MD simulation studies complement molecular docking studies, as MD simulations performed under biologically relevant environments give better insights into binding affinities of ligands, stability of overall systems, and better sampling of protein-ligand interactions [53]. The analysis of the evolution of protein structure from its initial equilibrated structure, as well as the evolution of different conformations of ligands in complex with protein, can be achieved through the measurement of RMSD, RMSF, and Rg. Here, the RMSD in protein backbone atoms below 0.2 nm indicates reasonable stability [54]. In the present study, AMP1, pfEMP1, and pfPK5 complexes have RMSD in backbone atoms around 0.2 nm, suggesting that the ligands have better affinity and stabilizing properties. The RMSD in ligand atoms typically showed that the ligands HC and HO have lower RMSD than CQ. HC and HO ligands have quite rigid structures with fewer rotatable bonds than CQ. Thus, in the case of CQ more conformations are possible and evidently, the conformations might be quite different from the initial equilibrated structure, which results in slightly higher RMSD values.

The RMSF in side chain atoms indicates mobile parts of the proteins under study. Lower RMSF indicates structural rigidity, while higher RMSF indicates structural mobility and conformational adaptation [55]. Both the chains of AMA1 are quite stable in complex with ligands. However, the ligand HC induces structural flexibility in AMA1. The RMSF in MSP1, pfCRT, and pfEMP1 represent the stability of the respective proteins in complex with the ligands. In the case of pfPK5, HC produced large structural changes, probably adversely affecting the stability.

The analysis of Rg points out the compactness of the system, where lower Rg represents a more compact structure [56]. In the case of AMA1, the complex with HC adversely affected the AMA1, resulting in destabilization after around 50 ns simulation period. The complexes of other proteins *viz.* MSP1, pfCRT, pfEMP1, and pfPK5 suggested reasonably compact structures.

The hydrogen bond analysis provides valuable insights into binding affinity as the more hydrogen bonds between ligand and protein, the better the binding affinity and corresponding stability of the resulting protein-ligand complex [57]. The contact frequency analysis provides valuable information about time-dependent key contacts between ligand and protein residues within a distance of 0.35 nm [58]. In the present work, the contact frequency was measured between the ligands and residues within 0.35 nm, which typically represents hydrogen bond distance. In the case of AMA1, only CQ showed reasons ligand chloroquine (CQ), Harman-3- carboxylic acid (HC), and Harmine N-oxide showed reasonably good binding free energies for the targets AMA1, MSP1, pfCRT, pfEMP1, and pfPK5. To get deeper insights into the binding modes of these ligands and to study the stabilization of these protein targets in complex with the ligands, 100 ns MD simulations were performed. Typically, MD simulation studies complement molecular docking studies, as MD simulations performed under biologically relevant environments give better insights into binding affinities of ligands, stability of overall systems, and better sampling of protein-ligand interactions [53]. The analysis of the evolution of protein structure from its initial equilibrated structure, as well as the evolution of different conformations of ligands in complex with protein, can be achieved through the measurement of RMSD, RMSF, and Rg. Here, the RMSD in protein backbone atoms below 0.2 nm indicates reasonable stability [54]. In the present study, AMP1, pfEMP1, and pfPK5 complexes have RMSD in backbone atoms around 0.2 nm, suggesting that the ligands have better affinity and stabilizing properties. The RMSD in ligand atoms typically showed that the ligands HC and HO have lower RMSD than CQ. HC and HO ligands have quite rigid structures with fewer rotatable bonds than CQ. Thus, in the case of CQ more conformations are possible and evidently, the conformations might be quite different from the initial equilibrated structure, which results in slightly higher RMSD values.

The RMSF in side chain atoms indicates mobile parts of the proteins under study. Lower RMSF indicates structural rigidity, while higher RMSF indicates structural mobility and conformational adaptation [55]. Both the chains of AMA1 are quite stable in complex with ligands. However, the ligand HC induces structural flexibility in AMA1. The RMSF in MSP1, pfCRT, and pfEMP1 represent the stability of the respective proteins in complex with the ligands. In the case of pfPK5, HC produced large structural changes, probably adversely affecting the stability.

The analysis of Rg points out the compactness of the system, where lower Rg represents a more compact structure [56] In the case of AMA1, the complex with HC adversely affected the AMA1, resulting in destabilization after around 50 ns simulation period. The complexes of other proteins *viz.* MSP1, pfCRT, pfEMP1, and pfPK5 suggested reasonably compact structures.

The hydrogen bond analysis provides valuable insights into binding affinity as the more hydrogen bonds between ligand and protein, the better the binding affinity and corresponding stability of the resulting protein-ligand complex [57]. The contact frequency analysis provides valuable information about time-dependent key contacts between ligand and protein residues within a distance of 0.35 nm [58]. In the present work, the contact frequency was measured between the ligands and residues within 0.35 nm, which typically represents hydrogen bond distance. In the case of AMA1, only CQ showed reasonably consistent hydrogen bonds and residues with higher contact frequencies. In the case of MSP1, the ligand HC has more consistent hydrogen bonds and residues Thr48 and Asn44 with over 75% contact frequency. The ligand CQ and HO has fewer hydrogen bonds and less contact frequency with the residues within 0.35 nm. The ligand HC might have better binding affinity than the ligand CQ and HO. While, in the case of pfCRT, the ligand HO showed a greater number of hydrogen bonds and the residues Asn246 and Arg231 with around 75% contact frequency, which suggests its better binding affinity than CQ and HC. In the case of pfEMP1, the ligands HC and HO having more hydrogen bonds and residues with over 75% contact frequency have better binding affinity than the CQ. However, in the case of pfPK5, though there are a greater number of frequent hydrogen bonds, the lower contact frequency in HC and HO suggests their weaker binding affinity.

Gibb’s free energy analysis of protein-ligand conformations from MD simulations provides valuable insights into the number of lowest energy and metastable conformations [59]. In the present study, Gibb’s free energy was analyzed for the protein-ligand conformations based on conformations isolated from essential dynamics. Here, two principal components were used as reaction coordinates. The results complement the hydrogen bond and contact frequency analysis. The AMA1 complex with CQ has more metastable conformations, suggesting better stability.

Similarly, the ligand HC, having more hydrogen bonds and residues with better contact frequency showed more metastable conformations. In the case of pfCRT, the complexes with CQ and HO showed more metastable conformations, suggesting better binding affinities of the ligands CQ and HO. In the case of pfEMP1, all the ligands have more metastable conformations, suggesting better binding affinities of these ligands. In the case of pfPK5, the complex of CQ only showed a small number of metastable conformations, while the ligands HC and HO almost lack metastable conformations, suggesting their poor binding affinities.

The MM-PBSA calculations on reasonably stable simulation trajectories provide probable binding free energy insights. However, in the present work, only the enthalpic energy contributions are calculated as relative binding free energies (ΔTOTAL) of respective ligands, ignoring the conformational entropic contributions upon protein-ligand binding [34]. The ligand CQ has better relative binding free energy against AMA1, MSP1, pfCRT, and pfPK5. However, the ligand HO has better relative binding free energy against pfEMP1. The ligand HC also has reasonably good and favorable relative binding free energy against pfEMP1 and pfCRT compared to CQ. Together with the results of hydrogen bond, contact frequency, and Gibb’s free energy analysis, the MM-PBSA analysis suggests comparably good binding affinity of compound HC and HO against pfCRT and pfEMP1. As these drug candidates showed relatively a significant result in *Insilco* analysis with having no-toxic or altered effect this harmala alkaloids could be considered as novel therapeutics. Further *in-vivo* and *in-vitro* study is recommended on these potential metabolites.

## 5. Figure Legends

Figure 1. Flowchart depicting the stepwise methods followed in the whole study.

Figure 2. RMSD, Rg, and RMSF analysis for AMP1 complexes.

Figure 3. Hydrogen bond and contact frequency analysis for AMP1 complexes.

Figure 4. RMSD, Rg, and RMSF analysis for MSP1 complexes.

Figure 5. Hydrogen bond and contact frequency analysis for MSP1 complexes.

Figure 6. RMSD, Rg, and RMSF analysis for pfCRT complexes.

Figure 7. Hydrogen bond and contact frequency analysis for pfCRT complexes.

Figure 8. RMSD, Rg, and RMSF analysis for pfEMP1 complexes.

Figure 9. Hydrogen bond and contact frequency analysis for pfEMP1 complexes.

Figure 10. RMSD, Rg, and RMSF analysis for pfPK5 complexes.

Figure 11. Hydrogen bond and contact frequency analysis for pfPK5 complexes.

Figure 12. Gibb’s free energy analysis. A) AMP1, B) MSP1, C) pfCRT, D) pfEMP1, and E) pfPK5.

## 6. Supplementary Information

Table S1: Functions of target proteins

Table S2: List of Harmala Alkaloid Plant metabolites containing anti-protozoal activity

Table S3: Binding Affinity of target proteins with enlisted metabolites

Figure S1: 3D visual representation of target proteins

Figure S2: Binding site of AMA1 protein with A) Harmalol B) Harmalol Chloride C) Harman- 3-Carboxylic acid and D) Harmin-N-oxide

Figure S3: Binding site of MSP1 protein with A) Harmalol B) Harmalol HydroChloride C) Harman-3-Carboxylic acid and D) Harmin-N-oxide

Figure S4: Binding site of pfCRT protein with A) Harmalol B) Harmalol HydroChloride C) Harman-3-Carboxylic acid and D) Harmin-N-oxide

Figure S5: Binding site of pfPK5 protein with A) Harmalol B) Harmalol HydroChloride C) Harman-3-Carboxylic acid and D) Harmin-N-oxide

Figure S6: Binding site of pfEMP1 protein with A) Harmalol B) Harmalol HydroChloride C) Harman-3-Carboxylic acid and D) Harmin-N-oxide

## Acknowledgements

The authors express their sincere gratitude for the computational support, resources, and collaborative assistance provided by the International Foundation for Collaborative Research (IFCR). As an international platform, IFCR is committed to equipping research and higher study enthusiasts with advanced research skills and publication opportunities. With administrative collaborations in numerous countries, especially in India, Bangladesh, and Africa, IFCR’s support has been crucial in the successful completion of this research.

