## Supplementary Table for "Exploring Harmala Alkaloids as Novel Antimalarial Agents against *Plasmodium falciparum* through Bioinformatics Approaches"

**Supplementary Tables**

**Table S1:** Functions of target proteins

| **Proteins** | **Functions** |
| --- | --- |
| PfCRT protein | PfCRT mutations may modify the chloroquine flow or decrease the drug's affinity for hematin.(Fidock et al., 2000)  An impact on the pH of the parasite's digestive vacuole.(Fidock et al., 2000)  Hypersensitivity To Other Antimalarials(Johnson et al., 2004)  Localizes to the membrane of the parasite's internal digestive vacuole (Lehane & Kirk, 2008)  PfCRT plays a role to degrade hemoglobin taken up by the parasite from its host cell(Sanchez et al., 2022).  Less is understood about PfCRT's non-drug related activity, but a recent study based on study done in a heterologous expression system suggests a role in oligopeptide transport.(Sanchez et al., 2022) |
| *P. falciparum* erythrocyte membrane protein 1 (PfEMP1) | PfEMP1 proteins enables parasites to evade host immunity and modifies parasite tropism for different microvascular beds.(Smith, 2014)  P. falciparum erythrocyte membrane protein 1 (PfEMP1) family mediates vascular endothelial binding (Smith, 2014)  PfEMP1 mostly control the virulence of parasite(Smith, 2014)  PfEMP1 enables the parasite to avoid splenic clearance (D'Ombrain et al., 2007)  PfEMP1 suppresses the production of the cytokine interferon-gamma by human peripheral blood mononuclear cells early after exposure to P. falciparum.(D'Ombrain et al., 2007)  PfEMP-1 is thought to interact with the CD36 receptor on antigen-presenting cells to control host immune responses(D'Ombrain et al., 2007). |
| Merozoite Surface Protein 1 (MSP1) | One of the most promising candidates for a vaccination against erythrocytic malaria parasites is merozoite surface protein-1(42) (MSP-1(42)).(Angov et al., 2003)  O-GlcNAc-modified MSP-1 N-terminal fragments tend to localise within the parasitophorous vacuolar membrane(Hoessli et al., 2003)  Antibodies to MSP1-19 may act as a marker of protective immunity.(Wilson et al., 2011)  It has been demonstrated that MSP1-19 binds to an RBC protein known as Band 3(Wilson et al., 2011)  MSP1-42 may interact with heparin-like molecules on RBCs, according to recent investigations(Wilson et al., 2011) |
| Apical Membrane Antigen 1 (AMA1) | AMA1 is a highly immunogenic protein(Healer et al., 2005)  Pfama1 Contributes to Erythrocyte Invasion via Specific Chimeric Ama1 Proteins That Include Pfama1 Domains I–III(Healer et al., 2005)  AMA1 is a significant genetic marker for investigating the relationships between malaria epidemiology and the adaptation of parasite populations (to the human host immune system).(Quang et al., 2009)  AMA1 forms a complex with parasite rhoptry neck (RON) proteins as part of the moving junction that develops between the host cell and the invading parasite.(A MacRaild et al., 2011)  It has been determined through structural analysis of AMA1 alone and in complexes with antibodies that prevent host cell invasion that the AMA1/RON complex is assembled in a manner that conserves a hydrophobic cleft.(A MacRaild et al., 2011) |
| *Plasmodium falciparum* protein Kinase 5 (PfPK5) | PfPK5 is necessary to activate or maintain the parasite S-phase.(Graeser et al., 1996)  In order to prevent cerebral malaria, P. falciparum Protein Kinase 5 (PfPK5) opens the prospect of attacking P. falciparum's life cycle specifically. A class of structurally conserved mammalian kinases called the cyclin-dependent kinases (CDKs) and PfPK5 share a significant degree of sequence similarity (>58%). The main regulatory components in charge of the mammalian cell cycle's orderly advancement are the CDKs.(Keenan & Welsh, 2004)  PfPK5 appears to have a part in the development of asexual blood stage parasites since it is expressed mostly during the ring stage of the life cycle of parasites.(Deng & Baker, 2002) |
